# Dora, a key component of target-directed miRNA degradation, is essential for local genomic amplification in *Drosophila* ovarian follicle cells

**DOI:** 10.1101/2025.02.20.639214

**Authors:** Natalia Akulenko, Oxana Olenkina, Elena Mikhaleva, Sofya Marfina, Anastasia Krylova, Stepan Toshchakov, Sergei Ryazansky

**Affiliations:** NRC “Kurchatov Institute”, Kurchatov sq. 2, Moscow 123182, Russia; Lomonosov Moscow State University, Faculty of Biology, Leninsky gory, 1/12, Moscow 125047, Russia

**Keywords:** *Drosophila*, oogenesis, Dora, TDMD, miRNAs, Ttk69, local genome amplification

## Abstract

The ubiquitin ligase receptor Dora, the *Drosophila* homolog of ZSWIM8, is a key component of the target-directed microRNA degradation (TDMD) pathway. Previous studies have implicated TDMD – and, consequently, ZSWIM8/Dora – in various developmental processes. Here, we investigate the role of Dora in *Drosophila* oogenesis, focusing on its function in ovarian somatic cells. We generated a fly strain with an endogenously tagged Dora protein and observed its presence in both germline and somatic follicular cells of the ovaries. Somatic knockdown of *dora* revealed its essential role in normal eggshell formation. Specifically, its loss led to reduced chorion and vitelline transcript levels and decreased chorion gene amplification, both of which are critical for eggshell protein production. Somatic depletion of Dora did not affect the abundance of other known regulators of eggshell formation, including Ttk69, Cut, miR-7 or miR-318 indicating that Dora functions independently of these pathways. Although a direct link between TDMD and chorion eggshell development remains to be confirmed, our findings clearly highlight Dora as a critical regulator in this process. These results pave the way for further investigation into the specific role of TDMD and provide new insights into the regulatory mechanisms underlying animal oogenesis.

## Introduction

Target-directed microRNA degradation (TDMD) is a sequence-specific miRNA turnover process in which animal miRNAs are degraded through non-canonical binding with specialized complementary RNAs, known as triggers (Akulenko et al., 2023; Ameres et al., 2010; Bitetti et al., 2018; Buck et al., 2010; Cazalla et al., 2010; Ghini et al., 2018; Han et al., 2020; Kingston et al., 2022; Kleaveland et al., 2018; Lee et al., 2013; Libri et al., 2012; Marcinowski et al., 2012; Sheng et al., 2023; Shi et al., 2020). In these duplexes, a miRNA pairs with a trigger along its entire length, except for a central region containing several mismatched nucleotides. In humans this duplex induces conformational changes in the associated Argonaute protein (Ago), allowing it to be recognized by the receptor ZSWIM8, a component of the Cullin-RING ligase (CRL3^TDMD^) complex. CRL3^TDMD^ in turn promotes Ago ubiquitination, leading to its degradation in proteasomes. The released miRNAs are further degraded (Han et al., 2020; Sheu-Gruttadauria et al., 2019; Shi et al., 2020). In addition to promoting miRNA decay, binding to TDMD triggers also induces trimming and tailing of the miRNA’s 3’-terminus (Ameres et al., 2010; Han et al., 2020; Shi et al., 2020). ZSWIM8 homologs, such as Dora in *Drosophila* and ebax-1 in nematodes, also play roles in TDMD (Akulenko et al., 2023; Donnelly et al., 2022; Kingston et al., 2022; Nahar et al., 2024; Shi et al., 2020; Stubna et al., 2024).

TDMD has been reported to be essential for proper embryonic development in both *Drosophila* and mice. In *Drosophila*, the loss of *dora* results in embryonic lethality and defects in cuticle formation, driven by the upregulation of miRNAs belonging to the miRNA-310 family (Kingston et al., 2022). Similarly, in mice, the loss of *Zswim8* leads to embryonic lethality accompanied by severe defects in heart and lung development as well as reduced body size (Jones et al., 2023; Shi et al., 2023). In the mammalian brain, the lncRNA *Cyrano* triggers TDMD of miR-7 leading to the accumulation of another miR-7 target *Cdr1as*, a circular RNA crucial for regulating neuronal activity (Kleaveland et al., 2018). Disruption of this pathway impairs neuronal functions. TDMD is also implicated in modulating animal behavior in zebrafish and mice. Specifically, the lncRNA *libra* in zebrafish and the mRNA *Nrep* in mice induce the degradation of miR-29b in certain brain cells, influencing exploratory and anxiety-related behaviors (Bitetti et al., 2018). Moreover, several herpesviruses exploit TDMD by expressing transcripts that induce the degradation of host miRNAs, including miR-17, miR-20a, miR-29, and miR-16, which would otherwise restrict viral replication (Buck et al., 2010; Cazalla et al., 2010; Lee et al., 2013; Libri et al., 2012; Marcinowski et al., 2012). In *Drosophila* ovarian somatic cell culture, *dora* knockdown inhibits the Notch signaling pathway (Akulenko et al., 2023). Additionally, degradation of human anti-apoptotic miR-221/222 through interaction with *BCL2L11* mRNA, which encodes the pro-apoptotic BIM protein, effectively enhances apoptosis. In this case the TDMD trigger and the encoded protein cooperate within the same pathway (Li et al., 2021). Notably, TDMD triggers are highly enriched in genes linked to carcinogenesis (Simeone et al., 2022) highlighting their potential role in cancer biology. However, the biological functions of the conserved TDMD pathway are likely much broader and remain largely understudied.

A promising model for studying the biological role of TDMD is *Drosophila* oogenesis, which is well-characterized, easy to study, and tightly regulated by miRNAs. *Drosophila* oogenesis involves several distinct stages [reviewed in (Berg et al., 2024)]. In early oogenesis (stages S1–S6), germline stem cells in the germarium differentiate into cysts, consisting of an interconnected oocyte and 15 nurse cells. Simultaneously, somatic stem cells give rise into several cell types, including follicle cells, which surround the germline cyst and mitotically proliferate. During mid-oogenesis (S7–S10A) the oocyte undergoes significant growth due to nutrient uptake and cytoplasmic transfer from nurse cells. Its nucleus migrates to the anterior dorsal corner, initiating dorsal-ventral patterning through Gurken signaling. Follicle cells undergo the M/E switch, transitioning from mitosis to genome replication without cell division (endocycles), leading to their increased ploidy. In late oogenesis (S10B–S14), nurse cells dump their remaining cytoplasmic content into the oocyte before undergoing apoptosis. The oocyte then undergoes partial meiosis, arresting at metaphase I to generate a haploid nucleus. Additionally, follicle cells transition from endocycles to a gene amplification program required for chorion gene expression (the E/A switch). These follicle cells secrete vitelline and chorion proteins to form eggshell layers and later specialize into eggshell structures such as dorsal appendages, the operculum, and the micropyle. The Notch and ecdysone signaling pathways, which interact and modulate each other’s activity, play multiple roles in oogenesis including cell fate determination, cell differentiation, nutrient uptake, regulation of the M/E and E/A switches, and eggshell maturation.

Previous studies have demonstrated that global disruption of the *Drosophila* miRNA system interferes with many processes in oogenesis (Azzam et al., 2012; Förstemann et al., 2005; Hatfield et al., 2005; Jiang et al., 2005; Jin and Xie, 2007; Park et al., 2007; Yang et al., 2007). Further research has identified specific miRNAs that regulate *Drosophila* oogenesis in both the germline (Hatfield et al., 2005; Nakahara et al., 2005; Shcherbata et al., 2007; Yu et al., 2009) and somatic cells (Ge et al., 2015; Huang et al., 2013; König and Shcherbata, 2015; Poulton et al., 2011; Sokol et al., 2008; Vilmos et al., 2013). These miRNAs regulate various processes, including stem cell proliferation and maintenance (Hatfield et al., 2005; Iovino et al., 2009; Jin and Xie, 2007; König and Shcherbata, 2015; Shcherbata et al., 2007; Sokol et al., 2008; Vilmos et al., 2013; Yu et al., 2009), cell fate determination (Poulton et al., 2011; Yatsenko and Shcherbata, 2018; Yoon et al., 2011), and border cells migration toward the oocyte (Kugler et al., 2013; Xia et al., 2024). The miRNA system in somatic cells has been reported to promote Notch signalling by repressing the Notch ligand Delta (Poulton et al., 2011) and the negative regulator Tom (Yatsenko and Shcherbata, 2018), as well as to modulate the ecdysone pathway (Xia et al., 2024). Certain miRNAs regulate anterior-posterior and dorsal-ventral patterning of egg chambers and are involved in eggshell maturation (Ge et al., 2015; Huang et al., 2013; Iovino et al., 2009).

Interestingly, miRNAs control eggshell maturation through two distinct mechanisms. The first one involves miR-7-dependent repression of the transcription factor Tramtrack69 (Ttk69) (Huang et al., 2013) having multiple roles in oogenesis. Ttk69 is a key transcriptional repressor coordinating developmental transitions in a follicular epithelium, including the M/E (Jordan et al., 2006; Sun and Deng, 2007) and E/A switches (Boyle and Berg, 2009; Huang et al., 2013; Sun and Deng, 2007), as well as dorsal appendage morphogenesis (Boyle et al., 2010; Boyle and Berg, 2009; French et al., 2003), and chorion gene expression (Boyle and Berg, 2009; French et al., 2003). Repression of *ttk69* by miR-7 is crucial for proper timing of the E/A switch in follicle cells during S10A/B stages (Huang et al., 2013).

The second mechanism involves miR-318, which acts independently but cooperatively with Ttk69 to regulate the progression of the E/A switch (Ge et al., 2015). Although loss of miR-318 does not prevent the initiation of the E/A switch, it impairs the subsequent gene amplification, which is further exacerbated by additional *ttk69* depletion. The involvement of miRNAs in multiple aspects of oogenesis underscores the need for precise regulation of their spatio-temporal expression, raising the question of whether TDMD contributes to this regulatory control.

The *Drosophila* ZSWIM8 ortholog Dora, which contains SWIM domain and extended intrinsically disordered regions (IDRs), participates in the TDMD pathway in embryos and embryonic S2 cell culture (Kingston et al., 2022; Sheng et al., 2023; Shi et al., 2020). Recently, we demonstrated that Dora functions in TDMD within ovarian somatic cell (OSC) culture (Akulenko et al., 2023). In that study, we endogenously tagged the *dora* gene with a hemagglutinin (HA) sequence within the OSC genome. Here, we investigated the biological role of Dora in *Drosophila* ovaries. Our results show that *dora* knockdown in ovarian somatic cells disrupts the E/A switch as well as both chorion gene amplification and expression. Eggs laid by females with somatic *dora* gene knockdown exhibit defective dorsal appendages, indicating impaired eggshells. Notably, Ttk69, Cut, miR-7 and miR-318 expression levels remain unaffected by Dora depletion, suggesting that Dora regulates the E/A switch and chorion maturation independently of the *ttk69*/miR-7 and miR-318 pathways.

## Results

### Tissue expression profile of Dora in *Drosophila*

To investigate Dora in *Drosophila*, we used Cas9-mediated scarless genome editing (Bier et al., 2018) to generate a fly strain carrying endogenously HA-tag *dora* allele (Figure 1A). As part of this strategy, a *dsRed* marker controlled by eye-specific enhancer was integrated into the *dora* 3’UTR to facilitate selection of the edited genomes. This marker was later removed using PiggyBac transposase to restore the native 3’UTR.

**Figure 1.**
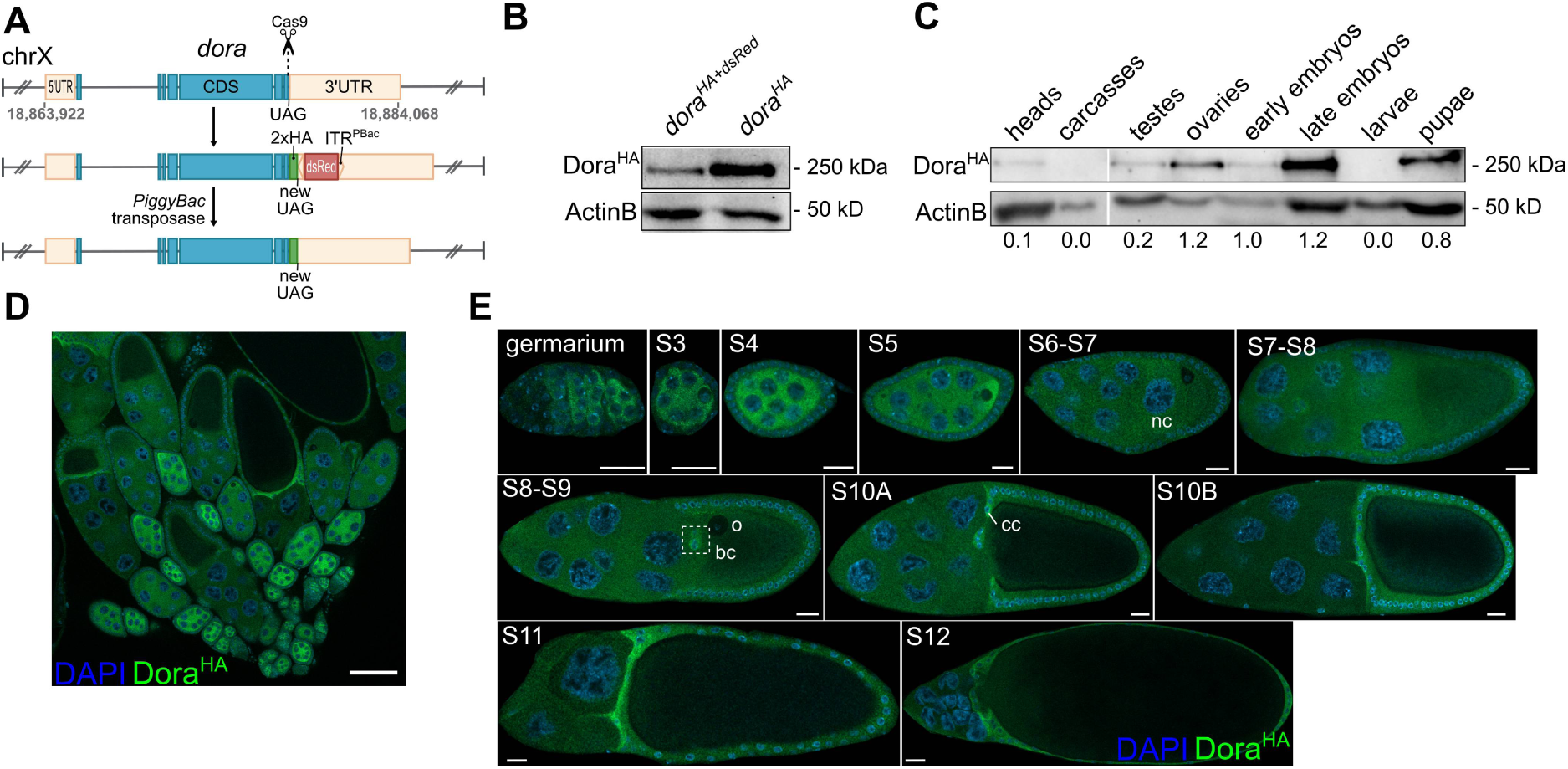
Endogenously tagged *dora* is ubiquitously expressed in *Drosophila* tissues. **A.** Schematic representation of the Cas9-mediated knock-in of an endogenous *dora* allele in the *Drosophila* genome. A 2xHA tag was inserted in frame at the C-terminus, while a *dsRed* marker was integrated into the *dora* 3’UTR. The *dsRed* cassette is flanked by inverted terminal repeats ITR of PiggyBac allowing its excision through PiggyBac transposase. **B**. Western blot of whole-body lysates using anti-HA antibodies. Samples include a strain carrying *dsRed* in the *dora^HA^*3’UTR, and a strain with *dsRed* excised. **C**. Tissue expression profile of Dora^HA^ was assessed by western blot with anti-HA antibodies. Probing with anti-actinB antibodies was used as a loading control in (**B**) and (**C**) panels. Ratios of HA to ActinB signals are shown below the panel. **D, E**. Immunostaining of *dora^HA^* ovaries using anti-HA antibodies (green). Nuclear DNA is counterstained with DAPI (blue). Cell types labeled include nurse cells (n), border cells (bc), maturing oocyte (o), centripetal cells (cc). Scale bars: 100 μm (**D**) and 20 μm (**E)**.

Western blot analysis using anti-HA antibodies showed that *dsRed* integration in the *dora* 3’UTR reduced *dora^HA^* expression whereas its excision restored expression level (Figure 1B). This indicates that the 3’UTR plays a critical role in regulating *dora* expression.

Next, we analyzed the tissue and developmental expression profile Dora^HA^. Samples were collected from early (2–12 h) and late (12–24 h) embryos, larvae, pupae, and adult flies, including dissected ovaries, testes, heads and carcass tissues. Western blot analysis revealed that Dora^HA^ was detected in nearly all examined samples, except larvae, where no expression was observed (Figure 1C). Notably, the Dora^HA^ levels were slightly elevated in ovaries and late embryos. These findings suggest that Dora and the TDMD pathway are broadly present across *Drosophila* tissues, while their function in larvae remain unclear.

### *Dora* is expressed in ovarian somatic cells

The tissue expression profile indicates that Dora^HA^ is highly abundant in late-stage embryos and ovaries (Figure 1C). Previous studies have shown that TDMD in embryos contributes to cuticle formation (Kingston et al., 2022; Sheng et al., 2023). Here, we focused on Dora’s function in ovaries.

Immunostaining of ovaries with anti-HA antibodies revealed *dora^HA^* expression in germline cells up to stages S5–S6, including stem cells in the germarium and the early oocyte, where Dora is asymmetrically localized around the nucleus (Figure 1D,E). Additionally, a strong Dora^HA^ signal was observed in various somatic cell types at S9–S12 (Figure 1E), particularly in border cells, centripetal cells, and epithelial follicular cells.

Border cells arise at S9 at the anterior tip of the egg chamber and migrate through the mass of nurse cells to their posterior border adjacent to the growing oocyte. These cells eventually differentiate into the micropyle of the laid egg (Montell, 2003; Saadin and Starz-Gaiano, 2016; Wu et al., 2008). Centripetal cells, which originate from somatic follicular cells covering the oocyte, specialize at S10A and migrate inward along the boundary between the oocyte and nurse cells, later forming the eggshell operculum (Wu et al., 2008). Follicular cells covering the maturing oocyte are essential for formation of chorion and vitelline layers in the eggshell (Wu et al., 2008).

We noticed that the expression domain of *dora^HA^* in ovarian somatic cells closely resembles that of the *cut* and *slbo* genes, which encode transcription factors with overlapping tissue expression patterns. *Cut* is expressed in follicular cells up to S6 and in border, centripetal, and follicular cells after S9–S10 (Jackson and Blochlinger, 1997; Levine et al., 2010; Sun and Deng, 2005) (Figure S1A). Similarly, *slbo* encodes the transcriptional factor C/EBP, which is expressed in border and centripetal cells and regulates their migration (Levine et al., 2010; Montell et al., 1992; Rørth et al., 2000; Saadin and Starz-Gaiano, 2016) (Figure S1B). Given the overlapping expressional domains of *slbo*, *cut*, and *dora^HA^*, we investigated whether Cut and Slbo might regulate *dora^HA^*. *In silico* analysis predicted multiple Cut and Slbo binding motifs for in the *dora* promoter region (motifs MA0244.1 and MA0188.1, respectively, from JASPAR (Rauluseviciute et al., 2024), p ≤ 10e-3) (Figure S2). To further test this hypothesis, we examined how depletion of these transcription factors affects Dora^HA^ abundance.

Neither the *slbo^01310^* mutation (Figure 2B’,B’’,C’,C’’) nor somatic *cut* knockdown affected Dora^HA^ distribution in ovaries (Figure 2D’,D’’). Western blot analysis confirmed that *cut* knockdown did not alter Dora^HA^ abundance (Figure 2D). Thus, we conclude that Slbo and Cut do not regulate the *dora* expression in ovarian somatic cells.

**Figure 2.**
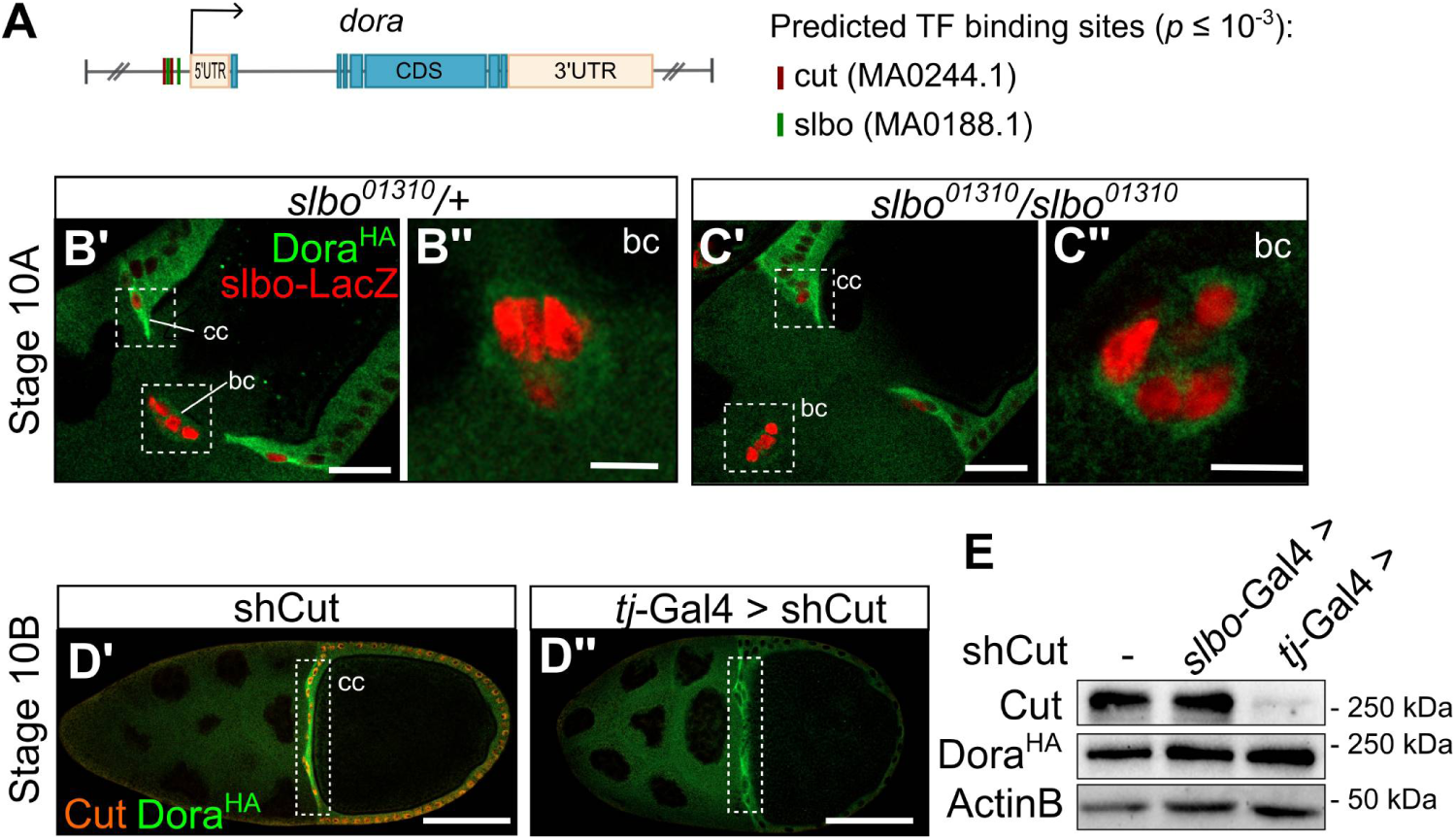
Expression of *dora^HA^*in ovarian somatic cells is independent of cut and *slbo*. **A.** Predicted binding sites of Cut and Slbo in the *dora* promoter region (-5 kb to +5 kb relative to the transcription start site). Binding sites were identified with FIMO (Grant et al., 2011) and JASPAR MA0244.1 and MA0188.1 weight matrices for Cut and Slbo respectively (p ≤ 10e-3). **B’–C’’.** Immunostaining of *dora^HA^* ovaries carrying *slbo^01310^* mutation using anti-HA antibodies (green). The *slbo^01310^* mutation results from the integration of a *lacZ* reporter construct into the *slbo* open reading frame; thus, *lacZ* expression is controlled by the *slbo* promoter and can be detected with anti-LacZ antibodies (red). **B’** and **C’** show S10A egg chamber highlighting oocyte (o), centripetal cells (cc), and border cells (bc). **B’’** and **C’’** are magnified views of border cells. Scale bars: 30 μm (B’, C’) and 10 μm (B’’, C”). **D’** and **D’’.** Immunostaining of Dora^HA^ with anti-HA antibodies (green) and Cut with anti-Cut antibodies (orange) in ovaries carrying *cut* somatic knockdown (*tj*-Gal4 > UAS-shCut). Scale bars: 100 μm (D’ and D’’). **E.** Western blot analysis of Dora^HA^ and Cut in ovaries with *cut* knockdown in all somatic cells (*tj*-Gal4 > UAS-shCut) or specifically in *slbo*-expressing somatic cells (*slbo*-Gal4 > UAS-shCut).

### Lack of follicular Dora impairs egg development

To further investigate the role of Dora in ovaries, we knocked down *dora^HA^* in ovarian somatic cells using the *tj-*Gal4 driver to induce UAS-shDora transgene. Ovarian chambers with Dora^HA^ depletion exhibited a complete loss of the anti-HA signal in somatic cells (Figure 3A), demonstrating the high efficiency of *dora* knockdown. Quantification of hatched eggs laid by females with depleted Dora revealed that nearly all eggs failed to develop into larvae (Figure 3B). Additionally, approximately half of the eggs displayed truncated dorsal appendages (Figure 3C). However, females in which *dora* was knocked down only in *slbo*-expressing border, centripetal, and polar cells showed normal egg hatching rates (Figure 3B) and displayed wild-type dorsal appendages. These findings indicate that Dora activity is required in epithelial follicular cells for normal egg development but is dispensable in *slbo*-expressing somatic cells.

**Figure 3.**
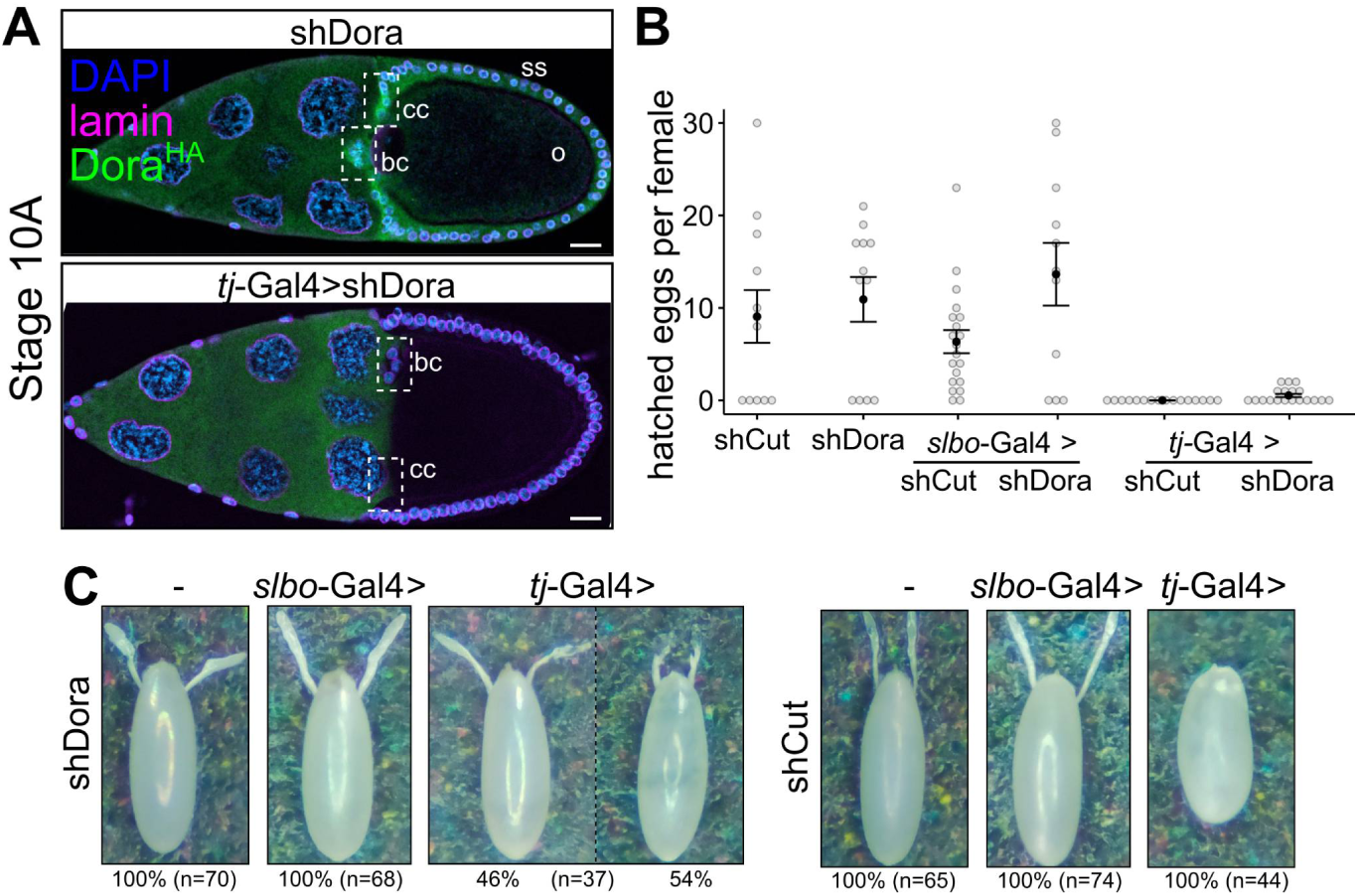
Knockdown of *dora* in ovarian follicular cells impairs egg development. **A.** Immunostaining of Dora^HA^ with anti-HA antibodies (green) and lamin with anti-lamin antibodies (purple) in ovaries where UAS-shDora is expressed in somatic cells under the control of the *tj*-Gal4 driver. Nuclear DNA is counterstained with DAPI (blue). Cell types labeled include nurse cells (n), border cells (bc), maturing oocyte (o), centripetal cells (cc), sheet follicular cells (ss). Scale bar: 50 μm. **B**. Number of hatched eggs laid by individual females of the specified genotypes. Egg hatching was assessed by the crossing of single females with 3–5 males followed by counting the number of hatched larvae. **C.** Microphotographs of eggs laid by females of the specified genotypes.

To gain further insight into Dora’s function, we compared the effects of its depletion with those of *cut* depletion. Cut is a homeodomain transcription factor that plays several essential roles in follicular cells during oogenesis, including promoting their proliferation during early oogenesis (Sun and Deng, 2005), ensuring proper differentiation during mid- and late oogenesis (Knapp et al., 2019; López-Schier and Johnston, 2001; Sun and Deng, 2005), and maintaining the functionality of border and centripetal cells (Levine et al., 2010). *Cut* is expressed exclusively in somatic tissues during the early and late oogenesis (Figure S1A).

Depletion of Cut resulted in abnormal egg chambers, ventralized eggs lacking dorsal appendages (Figure 3B), and a complete failure of egg hatching (Figure 3C). In contrast, the phenotypic effects of *dora* knockdown on egg maturation were less severe leading only to truncated appendages. This suggests that the defects caused by Dora depletion likely arise from a limited number of affected cells during late oogenesis, rather than a widespread disruption of somatic cell function.

### Dora is required for local replication and expression of chorion genes

Dorsal appendages are eggshell structures composed of differentiated follicular epithelial cells and chorion proteins. The developmental disruption of appendages, such as that observed in flies with somatic knockdown of *dora* and *cut* (Figure 3C), typically indicates a defective chorionic eggshell or disrupted oocyte dorsal-ventral (D/V) patterning. The truncated appendages in *dora* deficient eggs are unlikely to result from D/V patterning defects as the eggs were not ventralized. Instead, it is more likely that follicular cells in Dora-depleted ovaries failed to form a proper chorionic eggshell.

Eggshell formation is a multistep process that begins at S8–S9 with the expression of vitelline genes (Cavaliere et al., 2008). These genes encode vitelline proteins, which are secreted to form the first eggshell layer between an oocyte and follicular cells. At S10 follicular somatic cells transition from whole-genome endoreplication to local replication (amplification) of specific genomic regions containing chorion and vitelline gene clusters, a process known as the E/A switch (Hua and Orr-Weaver, 2017). Six known locally amplified genomic regions, referred to as <u>D</u>rosophila <u>a</u>mplicons in follicle <u>c</u>ells (DAFCs), exhibit varying levels of amplification. For instance, the DAFC-66D and DAFC-7F loci undergo 60–80-fold and 12–18-fold amplification, respectively (Hua and Orr-Weaver, 2017). Once amplification is completed, chorion gene transcription is activated, and the resulting proteins are secreted by follicular cells to form a multilayer eggshell envelope above the vitelline sheath.

We hypothesized that the failure of dorsal appendages formation was due to impaired genomic replication and/or transcription of chorion genes in follicular cells with *dora* knockdown. To test this, we measured expression levels of chorion and vitelline genes from the DAFC-66D and DAFC-7F loci using qRT-PCR. Ovaries with somatic *dora* knockdown exhibited reduced mRNA levels for all tested genes (Figure 4A). Additionally, qPCR analysis of genomic DNA revealed diminished genomic amplification at these loci (Figure 4B). This finding was further corroborated by genome-wide DNA sequencing (DNA-seq), which showed reduced read coverage at DAFC loci in Dora-depleted ovaries (Figure 4C, Figure S2).

**Figure 4.**
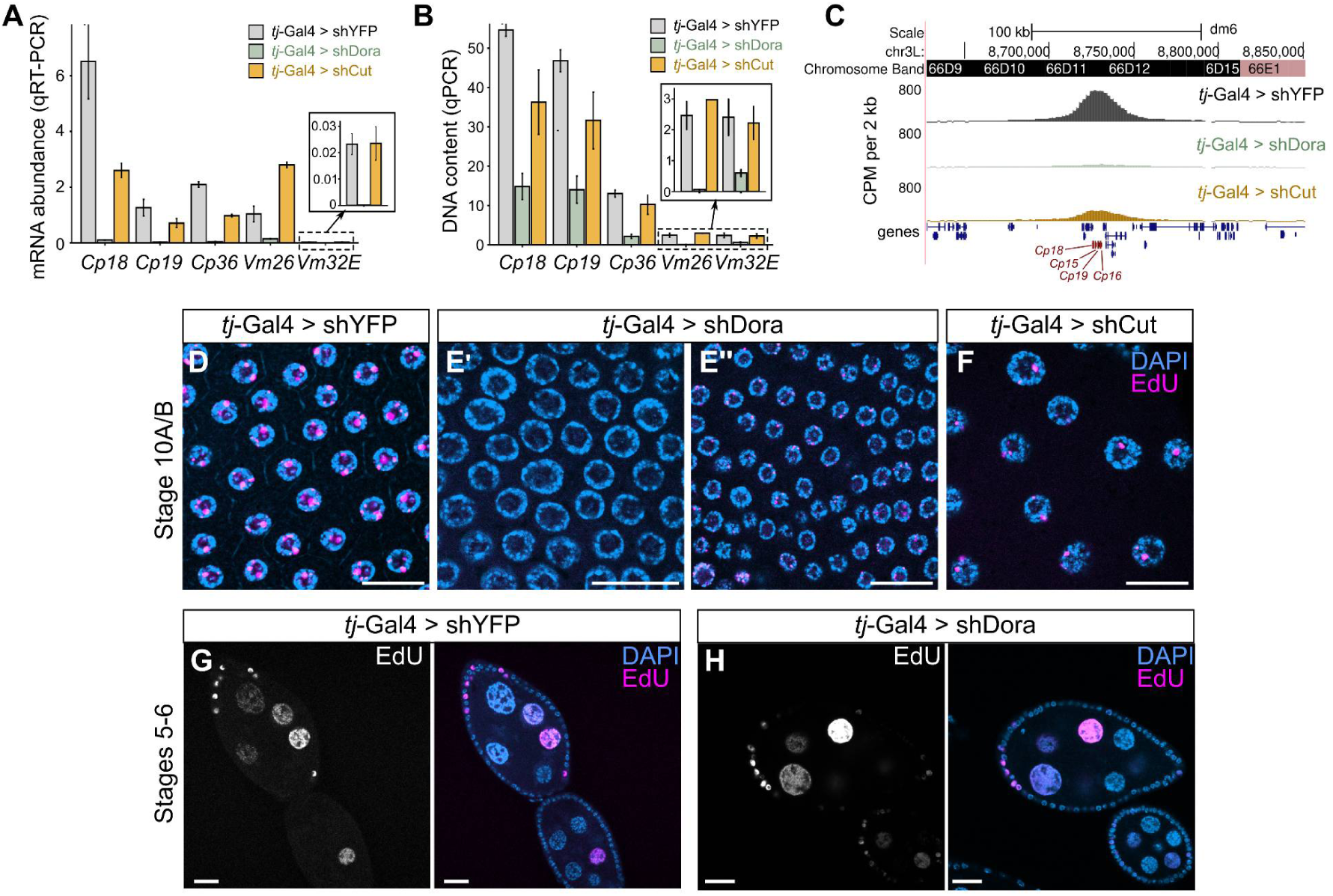
Somatic knockdown of *dora* reduces local replication and expression of chorion genes. **A**, **B**. Relative mRNA abundance (**A**) and DNA content (**B**) of chorion and vitelline genes in ovaries with somatic depletion of Dora^HA^ (*tj*-Gal4 > shDora) or Cut (*tj*-Gal4 > shCut) compared to the control strain (*tj-*Gal4 > shYFP). mRNA levels. mRNA abundance (A) was determined by qRT-PCR and normalized to *rp49*, while DNA content (B) was quantified by qPCR using total genomic DNA extracted from ovaries and normalized to the genomic region 60D. **C.** Normalized DNA-seq read coverage (CPM) at the DAFC-66D locus in ovaries with the specified genotype. **D–H.** EdU labeling assay (purple) in ovaries with the specified genotype. Nuclear DNA is counterstained with DAPI (blue). Follicular somatic cells in S10A/B (D, E’, E’’, F) and S5–S6 (G, H) are shown. Scale bars: 20 μm (D–H).

A reduction in genomic replication should correspond to decreased incorporation level of the nucleotide analog EdU into replicating DNA. In wild-type epithelial follicular cells at S10A/B, nuclei display 4–6 microscopically visible EdU incorporation spots, representing individual DAFCs (Figure 4D). However, in follicular cells with Dora depletion, most nuclei lacked visible EdU spots at this stage (Figure 4E’). In approximately 20% of ovarioles we observed small patches of cells displaying only one or two EdU spots (Figure 4E’’). These results demonstrate that Dora is essential for both amplification and transcription of chorion genes, processes critical for proper chorionic eggshell formation and dorsal appendage development.

Previous studies suggested that *cut* is not required for local genomic replication (Sun et al., 2008; Sun and Deng, 2005). However, we observed that ovaries with somatically depleted *cut* exhibited reduced gene amplification (Figure 4B,C, Figure S2), though the reduction was less pronounced compared to *dora* knockdown. Additionally, *cut* knockdown led to a decrease in the number and density of follicular cells at S10A/B, whose nuclei displayed fewer EdU spots (Figure 4F). Cut depletion also selectively reduced the abundance of chorion transcripts, but did not affect vitelline gene expression (Figure 4A). Thus, contrary to the previous reports (Sun et al., 2008; Sun and Deng, 2005), Сut may play a partial role in controlling chorion gene amplification and expression. This finding provides new insights into potential functions of Cut and warrants further investigation.

Importantly, Dora exerts a stronger influence on chorion gene amplification and expression, while the phenotypic effects of Dora depletion on eggs are less severe than those of *cut*. This suggests that Dora and Cut independently on each other regulate eggshell development through distinct mechanisms.

The local replication of specific genomic loci in S10A/B somatic cells is preceded by whole-genome endoreplication without cell divisions. Endoreplication begins at S5–S6 and continues until S10A/B leading to the polyploidization of follicular cells. To rule out the possibility that the failure of local replication upon Dora depletion was due to a general collapse of the replication machinery in the ovarian soma, we assessed endoreplication and polyploidization levels in these cells.

In wild-type ovaries, follicular nuclei in S5–S6 cells were uniformly labeled by EdU, consistent with whole-genome endoreplication (Figure 4G). Similarly, nuclei of somatic cells with Dora depletion displayed EdU labeling levels comparable to those in normal ovaries (Figure 4H). These findings demonstrate that endoreplication is unaffected by Dora depletion, highlighting its specific role in regulating localized genomic replication in S10A/B follicular cells.

### Dora is not required for *ttk69* expression

The developmental transitions of ovarian somatic cells from mitotic divisions to endoreplication at S5–S6 and from endoreplication to local replication at S10A/B are regulated by the Notch signaling pathway (Huang et al., 2013; Sun et al., 2008). Notch activity is essential for endoreplication, but its deactivation is required for the S10B gene amplification switch. Inactivation of the Notch pathway allows ecdysone signaling to induce *ttk69* expression (Sun et al., 2008), which in turn upregulates *cut* expression and suppresses genome-wide endoreplication, except at DAFC loci (Huang et al., 2013; Sun et al., 2008).

Given that Dora and Cut regulate eggshell development independently, we hypothesized that Dora does not regulate *ttk69*. To test this, we determined *ttk69* and *cut* expression domains by immunostaining in Dora-depleted ovaries. Our results confirmed that Dora depletion did not affect the abundance of either Ttk69 or Cut in S10A/B follicular cells and during early oogenesis (Figure 5).

**Figure 5.**
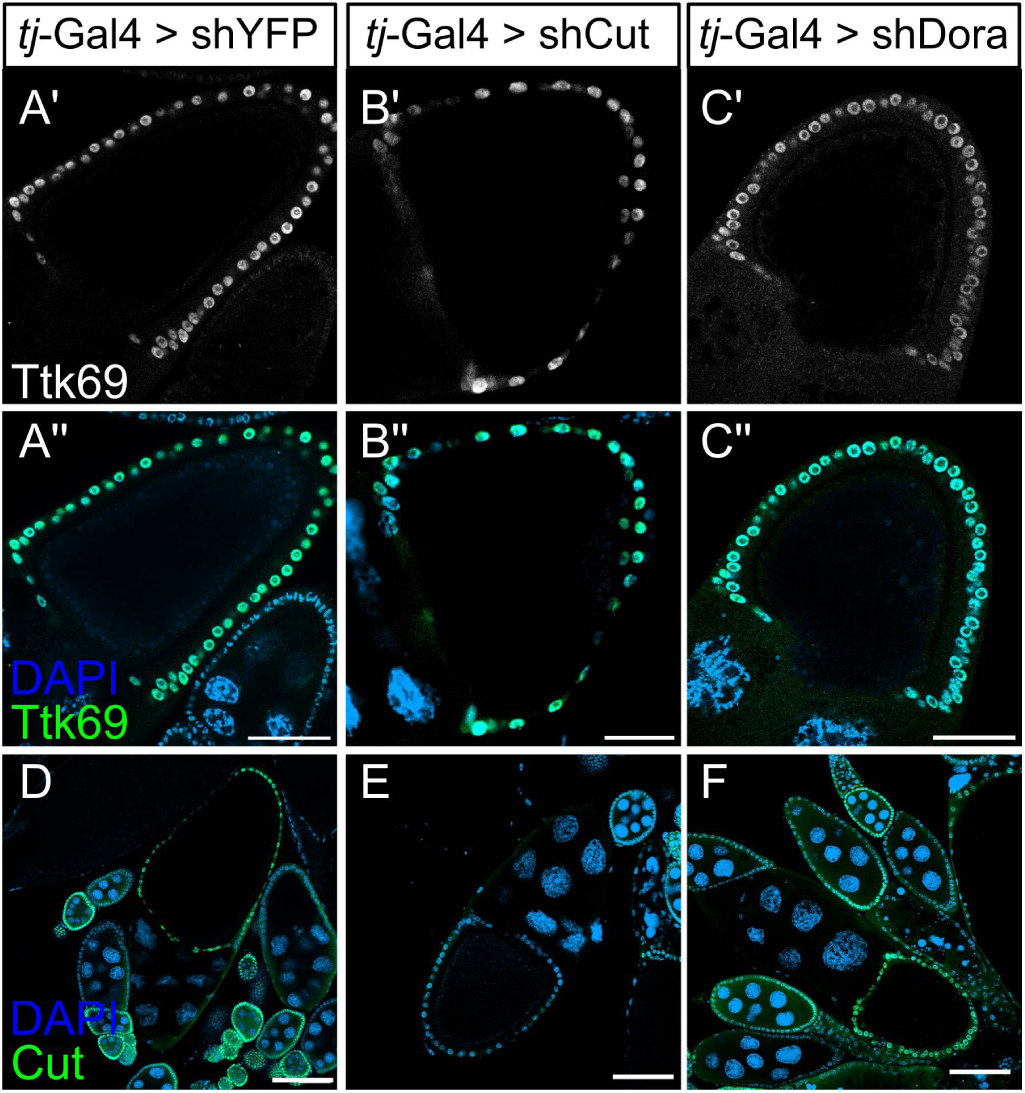
*ttk69* and *cut* expressions are not dependent on Dora. Immunostaining of S10A/B ovarian chambers with somatic depletion of Dora (*tj*-Gal4 > shDora) or Cut (*tj*-Gal4 > shCut) compared to the control strain (*tj*-Gal4 > shYFP) with anti-ttkC (**A–C**) or anti-Cut (**D–F**) antibodies. Nuclear DNA is stained with DAPI (blue). Scale bars: 50 μm (A–C) and 100 μm (D–F).

### Dora does not regulate miR-7 and miR-318 in ovarian somatic cells

Eggshell maturation is regulated by miRNAs through two distinct and independent mechanisms. The first involves miR-7-dependent repression of the transcription factor Ttk69, and the release of this repression at S10A/B is essential for the proper timing of the E/A switch (Huang et al., 2013). In the second mechanism, miR-318 functions after E/A initiation regulating the subsequent gene amplification process (Ge et al., 2015).

*Dora* encodes ZSWIM8 ortholog, which as part of the ubiquitin-ligase CRL3^TDMD^ complex, is involved in TDMD. Studies of its depletion in cell cultures (Akulenko et al., 2023; Kingston and Bartel, 2021; Sheng et al., 2023; Shi et al., 2020) and embryos (Kingston et al., 2022) have identified numerous miRNAs regulated by TDMD. Although Dora does not regulate *ttk69* expression and, therefore, miR-7 levels, it could still be involved in the TDMD-mediated degradation of miR-318 in ovarian somatic tissues.

To determine whether Dora regulates miR-318, we conducted high-throughput sequencing of ovarian small RNAs. We found that somatic knockdown of *dora* led to the statistically significant upregulation of only six canonical guide miRNAs (fold change ≥ 2, *p_adj_*≤ 10e-5) (Figure 6A). The number of *dora*-sensitive miRNAs was notably smaller compared to previously reported cell cultures and tissues in *Drosophila* and mammals (Akulenko et al., 2023; Kingston et al., 2022; Kingston and Bartel, 2021; Sheng et al., 2023; Shi et al., 2020).

**Figure 6.**
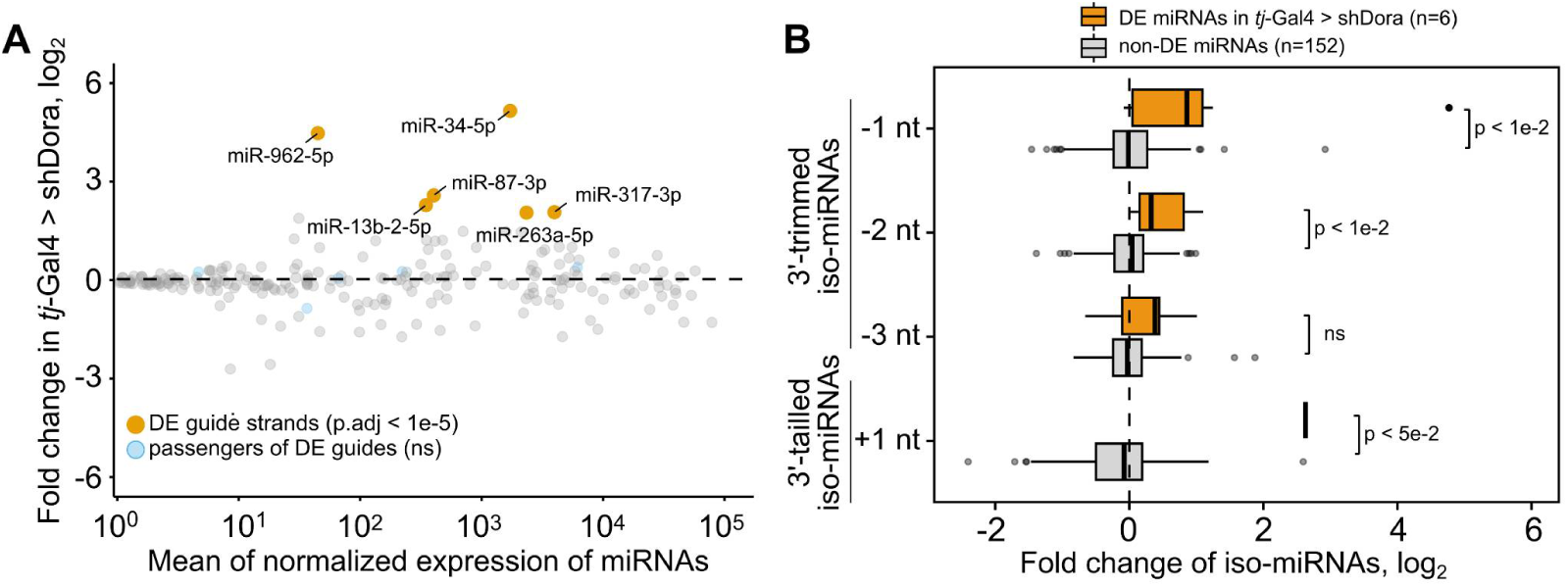
Differential expression of miRNAs upon Dora depletion. **A**. Fold-change expression of canonical miRNA forms in ovaries with somatic knockdown of *dora* (*tj*-Gal4 > shDora) compared to control ovaries (*tj*-Gal4 > shYFP) as determined by miRNA-seq (two biological replicates). Only guide strands (yellow dots) but not the corresponding passenger strands (light-blue dots) show differential expression. **B**. The barplot shows the fold-change expression of iso-miRNAs trimmed by one, two, or three nucleotides (-1, -2, or -3 nt) and tailed by one nucleotide (+1 nt) at the 3’-termini for upregulated miRNAs (orange bars) and miRNAs that do not change their expression levels (gray bars) upon somatic *dora* knockdown. There is only one Dora-regulated miRNA with accumulated 3’-tailed isoform, miR-87-3p. Statistical significance was assessed with the Wilcoxon unpaired test.w

The most upregulated miRNAs were miR-34-5p and miR-962-5p, while additional upregulated miRNAs included miR-87, miR-13b-2, miR-317, and miR-263a. The absence of miR-7 among them provides further evidence that Dora does not regulate the *ttk69*/miR-7 pathway, while the absence of miR-318 suggests that Dora may not contribute to E/A progression through miR-318. In agreement with previous reports (Akulenko et al., 2023; Han et al., 2020; Shi et al., 2020), only guide but not passenger strands were upregulated. We also detected the accumulation of tailing and trimming events of differentially expressed miRNAs on their 3’-termini (Figure 6B), reflecting the target-dependent trimming and tailing process associated with TDMD (Han et al., 2020; Shi et al., 2020). In general, the upregulation of miRNAs by *dora* loss likely indicates its involvement in TDMD in ovaries.

## Discussion

In this study, we demonstrated that Dora is highly expressed in ovarian somatic tissues and acts as a novel regulator of E/A switch initiation, as well as chorion and vitelline gene amplification and expression (Figure 4). It remains unclear whether the decrease in chorion gene expression observed in Dora-depleted ovaries is entirely due to their reduced genomic amplification (Cavaliere et al., 2008; Claycomb and Orr-Weaver, 2005; Hua and Orr-Weaver, 2017; Papantonis et al., 2015) or whether Dora independently regulates both expression and amplification.

Dora and its ZSWIM8 homologs are key components of TDMD, a process during which miRNAs are degraded upon complementary binding with specialized RNA triggers (Akulenko et al., 2023; Donnelly et al., 2022; Han et al., 2020; Kingston et al., 2022; Nahar et al., 2024; Shi et al., 2020; Stubna et al., 2024). While our study did not establish a direct link between TDMD and chorionic eggshell development, it raises the question of whether Dora functions in late oogenesis through miRNA inhibition.

Two miRNAs are known to regulate the initiation and progression of the E/A switch, miR-7 and miR-318 (Ge et al., 2015; Huang et al., 2013). Suppression of miR-7 in S10A/B somatic cells is necessary for activation of its target, the transcription factor Ttk69 (Huang et al., 2013), which in turn coordinates the E/A switch (Boyle and Berg, 2009; Huang et al., 2013; Sun and Deng, 2007) and chorion gene expression (Boyle and Berg, 2009; French et al., 2003). Ttk69 also plays a role in dorsal appendage morphogenesis (Boyle et al., 2010; Boyle and Berg, 2009; French et al., 2003). One possible hypothesis is that miR-7 suppression in S10A/B, and hence *ttk69* derepression, is determined by TDMD with Dora participation. This is supported by the fact that miR-7 is a conserved TDMD substrate in various animal tissues and cell cultures, including *Drosophila* ovarian somatic cell culture (Akulenko et al., 2023; Shi et al., 2023, 2020). However, *dora* depletion did not result in miR-7 accumulation (Figure 6), and Ttk69 and its positively regulated Cut protein were expressed as in normal ovaries (Figure 5). Thus, Dora controls the E/A switch and chorion amplification independently of the *ttk69*/miR-7.

Another possibility is that Dora determines the spatio-temporal expression domain of miR-318, which is required for gene amplification maintenance (Ge et al., 2015). However, we did not observe miR-318 accumulation in *dora*-depleted ovaries (Figure 6). Moreover, miR-318 is not required for E/A initiation but only for the progression of local amplification. This is evident from the fact that loss of miR-318 in S10A/B follicular cells only reduces but does not abolish the number of locally replicating regions (Ge et al., 2015). In contrast, Dora depleted follicular cells almost completely lacked these regions (Figure 4). These findings suggest that Dora acts earlier than miR-318 and is not involved in the miR-318-mediated pathway.

Taken together, our results demonstrate that Dora regulates chorion eggshell formation independently of miR-7 and miR-318 activity. However, six other Dora-regulated miRNAs (Figure 6) may contribute to this process. It is important to note that while our study focused on miRNAs regulated by Dora in ovarian somatic cells, miRNA-seq was performed on total ovaries which include both somatic and germline cell types. Therefore, some differentially expressed and low-abundance miRNAs in somatic cells may not have been detected due to their masking by larger germline miRNA pool. This suggests that our finding of six Dora-regulated miRNAs represents a conservative estimate of the total number involved in ovarian somatic regulation.

Among the identified miRNAs, miR-34 has multiple functions in *Drosophila* (Wang et al., 2022), including the regulation of border cell migration (Xia et al., 2024). Although miR-34 overexpression in border cells represses the *Eip74EF* gene target, which is required for their movement (Xia et al., 2024), we did not observe significant defects in border migration in Dora-depleted ovaries. This suggests that in our study, miR-34 was likely upregulated in other ovarian cell types rather than border cells. Two other Dora-regulated miRNAs, miR-13b-2 and miR-87, are maternally inherited (Chen et al., 2014; Marco, 2015, 2014). Loss of maternal and zygotic miR-87 expression leads to embryo lethality (Chen et al., 2014). Future studies should highlight the potential role of these and other Dora-regulated miRNAs (miR-962, miR-317, and miR-263a) in chorion gene amplification and expression.

Additionally, beyond their specific functions it will be valuable to determine whether Dora mediates the spatiotemporal regulation of these miRNAs in ovarian somatic cells. This could provide insights into how TDMD contributes to cell-type specific miRNA expression.

An alternative possibility is that Dora regulates chorion development through a miRNA-independent mechanism. While little is known about non-TDMD functions of ZSWIM8, it has been reported that ZSWIM8 acts as a quality control protein targeting misfolded Dab1 for degradation (Wang et al., 2023). Dab1 is essential for mammalian brain development, and loss of ZSWIM8 in mice led to defective brain cells and impaired learning and memory. The proposed mechanism suggests that ZSWIM8 recognizes Dab1 through its extended IDRs triggering its proteasomal degradation (Wang et al., 2023). It is possible that Dora also functions as a quality control factor for proteins essential to chorion gene amplification, which warrants further investigation.

Several other open questions about Dora remain. One is how Dora activity in late oogenesis is integrated with the Notch and ecdysone signaling pathways. The E/A switch is only possible through the coordinated action of these two pathways where Notch downregulation allows EcR to promote *ttk69* expression (Jordan et al., 2006; Sun et al., 2008). Our data indicate that Dora does not modulate the Notch pathway since *ttk69* expression remained unchanged in Dora-depleted ovaries.

However, it is unknown if Notch and ecdysone pathways may regulate *dora*. Additionally, our data suggest that Dora does not act upstream of Ttk69 and functions independently of Ttk69 positively regulated Cut. This raises another important question: does Dora act downstream or in parallel to Ttk69 in controlling chorion gene amplification? Understanding how Notch and ecdysone signaling influence Dora function and how Dora interacts with Ttk69 will be critical to fully elucidating underlying molecular mechanisms.

Interestingly, our findings of the role of Dora in chorion eggshell development parallel a recent report on TDMD’s involvement in embryonic cuticle formation (Kingston et al., 2022). This study found that the long ncRNA *Marge* acts as a TDMD trigger targeting six miRNAs from the miR-310 family for degradation, a process essential for normal cuticle formation. Without *Marge*-mediated degradation of miR-310 family members, the cuticle becomes enlarged, leading to embryonic lethality (Kingston et al., 2022). Thus, Dora plays a key regulatory role in both eggshell and cuticle formation utilizing distinct mechanisms throughout *Drosophila* development.

## Materials and Methods

### Data availability

Raw data from high-throughput sequencing of miRNAs and genomic DNA are available in NCBI BioProject (PRJNA1218501).

### Fly stocks

Flies were maintained on standard fly food and crossed at 25°C. The following strains from the Bloomington Drosophila Stock Center (BDSC; Bloomington, USA) were used: vasa-Cas9 (BDSC #51324), slbo^01310^ (BDSC #12227), piggyBac strain (BDSC #8285), UAS-shDora (BDSC #18553), UAS-Dcr2 (BDSC #24651), and CantonS (BDSC #64349). *tj*-Gal4 strain was from the Kyoto Drosophila Stock Center (#104055). Fly strains carrying UAS-shCut and UAS-shYFP transgenes were generated in this study. Transgenesis was performed using the phiC31 integration system and a fly strain with an attP landing site at 75A10 locus (BDSC #24862). Strains *slbo*-Gal4 (Novo16) (Ogienko et al., 2018) and slbo^01310^ were kindly provided by A. Ogienko.

### Egg hatching test

Fifteen 3-day-old females of the tested genotype were individually crossed with five 3-day-old wild-type CantonS males. Parents were removed from vials after five days and the number of hatched eggs from each cross was manually counted.

### Endogenous tagging of *dora*

Endogenous tagging of *dora* was performed using a scarless genome editing technique (Bier et al., 2018). A donor plasmid for homology-directed repair was constructed based on the pHD-2xHA-ScarlessDsRed vector (Addgene plasmid #80822; a gift from Kate O’Connor-Giles). This plasmid contained a 3xP3-dsRed marker flanked by the *dora* coding sequence in-frame with the 2xHA-tag sequence (left homology arm, LA) followed by the *dora* 3’UTR (right homology arm, RA). The RA was PCR-amplified from the vasa-Cas9 strain genome using primers carrying a mutated *Eco*RI site (Table S1) and inserted into the *Pst*I linearized pHD-2xHA-ScarlessDsRed vector using Gibson assembly (HiFi DNA Assembly, NEB). The LA was PCR-amplified from genomic DNA using primers carrying a mutated PAM site (Table S1) and introduced into *Eco*RI linearized pHD2xHA/Dora_RA plasmid using Gibson assembly. A guide (sgRNA) sequence was selected using TargetFinder (Gratz et al., 2014). Corresponding oligonucleotides (Table S1) were annealed and cloned into the pU6-BbsI-chiRNA vector (Addgene plasmid #45946; a gift from Melissa Harrison & Kate O’Connor-Giles & Jill Wildonger) linearized with *Bbs*I (Gratz et al., 2013). The donor plasmid pHD2xHA/Dora_RA&LA and pU6/sgRNA[Dora] were injected into *vasa*-Cas9 embryos. Founder males were individually crossed with *yw^67c23^* females to generate progeny carrying HA-tagged *dora* (marked by eye-expressed *dsRed*) while lacking the *vasa-Cas9* transgene (marked by germline-expressed *GFP*). To excise *dsRed* flanked by *piggyBac* termini from the *dora* 3’UTR, flies were crossed with a PiggyBac transposase expressing strain.

### Knockdown of *dora* and *cut*

To generate UAS-shCut and control UAS-shYFP expressing transgenes, corresponded oligonucleotides (Table S1) were synthesized, annealed, and cloned into the pWalium20 vector (DGRC #1472) linearized with *Nhe*I and *Eco*RI endonucleases (Ni et al., 2011). Transgenic fly strains were obtained using the phiC31 integration system (Bischof et al., 2007). Strain carrying UAS-shDora was obtained from BDSC (BDSC #18553). Knockdown was achieved by crossing UAS-shRNA strains and driver strains carrying UAS-Dcr2 and *tj*-Gal4 or *slbo*-Gal4.

### Western blot

Western blotting was performed on 6–7-day-old females, larvae, embryos and pupae as previously described (Akulenko et al., 2023). The following primary antibodies were used: mouse anti-HA (1:1,000, 6E2, Cell Signaling, #2367), mouse anti-Cut (1:200, DSHB, 2B10), and mouse anti-ActinB (1:1,000, Abcam, AB8224).

### Small RNA-seq

Small RNA fractions were extracted using RNAzol (MRC) from ovaries of 5-day-old flies. Libraries were prepared (two biological replicates of each genotype) as previously described (Akulenko et al., 2023) and subjected to high-throughput sequencing using NovaSeq 6000 (Illumina) at Novogen. Bioinformatic analysis was conducted as previously described (Akulenko et al., 2023).

### Genomic DNA-seq

Genomic DNA was extracted from ovaries of 5-day-old flies, ultrasonicated, and prepared for sequencing using NEBNext Ultra II DNA library preparation kit (NEB). Libraries were sequenced using NovaSeq 6000 (Illumina) at the Resource Center of the NRC “Kurchatov Institute”. A total of 11–15 million of paired-end reads (180–210-bp fragment sizes) were obtained, providing 12–19-fold euchromatic genome coverage. Reads were mapped to the *dm6* genome assembly using *bowtie2*, deduplicated, and single-mapped reads were counted in 2 kb bins.

### Immunostaining

Ovarian immunostaining was performed as previously described (Fefelova et al., 2022). The following primary antibodies were used: mouse anti-HA (1:300, 6E2, Cell Signaling, #2367), mouse anti-Cut (1:50, 2B10, DHSB), rat anti-TtkC (Khalisova et al., 2021), and mouse anti-LacZ (1:10, 40-1a, DSHB). Secondary Alexa Fluor antibodies (1:500, ThermoFisher Scientific) were used. Confocal microscopy was performed using a Zeiss LSM 510 META system and LSM 900 Confocal with an Airyscan2 super-resolution detector.

### EdU labeling assay

Ovaries from 5–7-day-old females were dissected and thoroughly combed in Schneider’s Medium supplemented with 10% fetal bovine serum. EdU (Invitrogen) with 20 μM final concentration was added, and samples were incubated at 25°C for 2 h. After incubation ovaries were washed with PBS, fixed in 4% formaldehyde for 20 min under rotation, and then washed twice with PBS. Fixed ovaries were permeabilized in PBTX (0.1% Tween20 and 0.3% Triton X-100) for 30 min and finally washed twice in PBS. Click chemistry reactions were carried out using Click-iT™ Cell Reaction Buffer Kit (Invitrogen) for 30 min at room temperature in the dark followed by PBS washing. Nuclei were counterstained with DAPI (1 μg/ml for 5 min) and ovaries were mounted on glass slides in SlowFade (Thermo Fisher Scientific) for confocal microscopy.

### qPCR

Transcript abundance of chorion and vitelline genes was measured by qRT-PCR as previously described (Akulenko et al., 2023) and normalized to *rp49* (2^-dCt^). Genomic DNA qPCR was performed using the same primers and normalized to the transcriptionally inert 60D region (2^-dCt^). Primers are listed in Table S1.

## Acknowledgments

We thank Yulii Shidlovsky (Institute of Gene Biology, Moscow, Russia) for the vasa-Cas9 fly strain; Anna Ogienko (Institute of Molecular and Cellular Biology, Novosibirsk, Russia) and Alexey Pindurin (Institute of Molecular and Cellular Biology, Novosibirsk, Russia; current affiliation is Annogen Bv, Amsterdam, Netherlands) for *slbo* fly strains; and Oxana Maksimenko (Institute of Gene Biology, Moscow, Russia) for providing anti-ttkC antibodies. We thank Leonid Kulakov for English editing. The work was carried out using a confocal microscope and NovaSeq 6000 at the Resource Center of the NRC “Kurchatov Institute”. The work was carried out within the state assignment of the NRC “Kurchatov Institute”.

## Author contributions

N.A. performed most experiments, including molecular cloning, western blotting, qRT-PCR, and library preparation for miRNA-seq and DNA-seq. O.O. handled fly maintenance, genetic crossings, and generation of transgenic strains. E.M. performed ovarian immunostaining. S.M. conducted molecular cloning, western blotting, and qRT-PCR. A.K. and S.T. performed DNA-seq. S.R. and N.A. conceived and designed the project and edited the manuscript; S.R. wrote the manuscript with input from N.A. All authors read and approved the final version.

---

**Figure S1.** Immunostaining of Cut (**A**) and Slbo (**B**) in ovaries of *CantonS* and *slbo^01310/^+* fly strains, respectively. Cell types labeled include border cells (bc), centripetal cells (cc), and polar cells (pc). Scale bar: 50 μm.

**Figure S2.** Normalized DNA-seq reads coverage (CPM) of locally replicated genomic loci of ovaries with somatic depletion of Dora (*tj*-Gal4 > shDora) or Cut (*tj*-Gal4 > shCut) compared to the control strain (*tj*-Gal4 > shYFP). A’, B’, C’ - DNA-seq read coverage along the entire chromosome 2L (A’), 3L (B’), and X (C’). A’’, A’’’, B’’, B’’’, C’’ - DNA-seq read coverage along the specified DAFC.

**Table S1.** Primers and oligonucleotides used for PCR, cloning, and miRNA-seq.

## References

Akulenko, N., Mikhaleva, E., Marfina, S., Kornyakov, D., Bobrov, V., Arapidi, G., Shender, V., Ryazansky, S., 2023. Evidence of target-mediated miRNA degradation in Drosophila ovarian cell culture. 10.1101/2023.08.30.555489

Ameres, S.L., Horwich, M.D., Hung, J.-H., Xu, J., Ghildiyal, M., Weng, Z., Zamore, P.D., 2010. Target RNA-Directed Trimming and Tailing of Small Silencing RNAs. Science 328, 1534–1539. 10.1126/science.1187058

Azzam, G., Smibert, P., Lai, E.C., Liu, J.-L., 2012. Drosophila Argonaute 1 and its miRNA biogenesis partners are required for oocyte formation and germline cell division. Developmental biology 365, 384–94. 10.1016/j.ydbio.2012.03.005

Berg, C., Sieber, M., Sun, J., 2024. Finishing the egg. GENETICS 226, iyad183. 10.1093/genetics/iyad183

Bier, E., Harrison, M.M., O’Connor-Giles, K.M., Wildonger, J., 2018. Advances in Engineering the Fly Genome with the CRISPR-Cas System. Genetics 208, 1–18. 10.1534/genetics.117.1113

Bischof, J., Maeda, R.K., Hediger, M., Karch, F., Basler, K., 2007. An optimized transgenesis system for Drosophila using germ-line-specific phiC31 integrases. Proceedings of the National Academy of Sciences of the United States of America 104, 3312–7. 10.1073/pnas.0611511104

Bitetti, A., Mallory, A.C., Golini, E., Carrieri, C., Carreño Gutiérrez, H., Perlas, E., Pérez-Rico, Y.A., Tocchini-Valentini, G.P., Enright, A.J., Norton, W.H.J., Mandillo, S., O’Carroll, D., Shkumatava, A., 2018. MicroRNA degradation by a conserved target RNA regulates animal behavior. Nature Structural & Molecular Biology 25, 244–251. 10.1038/s41594-018-0032-x

Boyle, M.J., Berg, C.A., 2009. Control in time and space: Tramtrack69 cooperates with Notch and Ecdysone to repress ectopic fate and shape changes during Drosophila egg chamber maturation. Development 136, 4187–4197. 10.1242/dev.042770

Boyle, M.J., French, R.L., Cosand, K.A., Dorman, J.B., Kiehart, D.P., Berg, C.A., 2010. Division of labor: Subsets of dorsal-appendage-forming cells control the shape of the entire tube. Developmental Biology 346, 68–79. 10.1016/j.ydbio.2010.07.018

Buck, A.H., Perot, J., Chisholm, M.A., Kumar, D.S., Tuddenham, L., Cognat, V., Marcinowski, L., Dölken, L., Pfeffer, S., 2010. Post-transcriptional regulation of miR-27 in murine cytomegalovirus infection. RNA (New York, N.Y.) 16, 307–15. 10.1261/rna.1819210

Cavaliere, V., Bernardi, F., Romani, P., Duchi, S., Gargiulo, G., 2008. Building up the Drosophila eggshell: First of all the eggshell genes must be transcribed. Developmental Dynamics 237, 2061–2072. 10.1002/dvdy.21625

Cazalla, D., Yario, T., Steitz, J.A., 2010. Down-Regulation of a Host MicroRNA by a Herpesvirus saimiri Noncoding RNA. Science 328, 1563–1566. 10.1126/science.1187197

Chen, Y.-W., Song, S., Weng, R., Verma, P., Kugler, J.-M., Buescher, M., Rouam, S., Cohen, S.M., 2014. Systematic Study of Drosophila MicroRNA Functions Using a Collection of Targeted Knockout Mutations. Developmental Cell 31, 784–800. 10.1016/j.devcel.2014.11.029

Claycomb, J., Orr-Weaver, T., 2005. Developmental gene amplification: insights into DNA replication and gene expression. Trends in Genetics 21, 149–162. 10.1016/j.tig.2005.01.009

Donnelly, B.F., Yang, B., Grimme, A.L., Vieux, K.-F., Liu, C.-Y., Zhou, L., McJunkin, K., 2022. The developmentally timed decay of an essential microRNA family is seed-sequence dependent. Cell Reports 40. 10.1016/j.celrep.2022.111154

Fefelova, E.A., Pleshakova, I.M., Mikhaleva, E.A., Pirogov, S.A., Poltorachenko, V.A., Abramov, Y.A., Romashin, D.D., Shatskikh, A.S., Blokh, R.S., Gvozdev, V.A., Klenov, M.S., 2022. Impaired function of rDNA transcription initiation machinery leads to derepression of ribosomal genes with insertions of R2 retrotransposon. Nucleic Acids Research 50, 867–884. 10.1093/nar/gkab1276

Förstemann, K., Tomari, Y., Du, T., Vagin, V.V., Denli, A.M., Bratu, D.P., Klattenhoff, C., Theurkauf, W.E., Zamore, P.D., 2005. Normal microRNA maturation and germ-line stem cell maintenance requires Loquacious, a double-stranded RNA-binding domain protein. PLoS biology 3, e236. 10.1371/journal.pbio.0030236

French, R.L., Cosand, K.A., Berg, C.A., 2003. The Drosophila Female Sterile Mutation twin peaks Is a Novel Allele of tramtrack and Reveals a Requirement for Ttk69 in Epithelial Morphogenesis. Developmental Biology 253, 18–35. 10.1006/dbio.2002.0856

Ge, W., Deng, Q., Guo, T., Hong, X., Kugler, J.-M., Yang, X., Cohen, S.M., 2015. Regulation of Pattern Formation and Gene Amplification During *Drosophila* Oogenesis by the miR-318 microRNA. Genetics 200, 255–265. 10.1534/genetics.115.174748

Ghini, F., Rubolino, C., Climent, M., Simeone, I., Marzi, M.J., Nicassio, F., 2018. Endogenous transcripts control miRNA levels and activity in mammalian cells by target-directed miRNA degradation. Nature Communications 9, 3119. 10.1038/s41467-018-05182-9

Grant, C.E., Bailey, T.L., Noble, W.S., 2011. FIMO: scanning for occurrences of a given motif. Bioinformatics 27, 1017–1018. 10.1093/bioinformatics/btr064

Gratz, S.J., Cummings, A.M., Nguyen, J.N., Hamm, D.C., Donohue, L.K., Harrison, M.M., Wildonger, J., O’Connor-Giles, K.M., 2013. Genome Engineering of *Drosophila* with the CRISPR RNA-Guided Cas9 Nuclease. Genetics 194, 1029–1035. 10.1534/genetics.113.152710

Gratz, S.J., Ukken, F.P., Rubinstein, C.D., Thiede, G., Donohue, L.K., Cummings, A.M., O’Connor-Giles, K.M., 2014. Highly Specific and Efficient CRISPR/Cas9-Catalyzed Homology-Directed Repair in Drosophila. Genetics 196, 961–971. 10.1534/genetics.113.160713

Han, J., LaVigne, C.A., Jones, B.T., Zhang, H., Gillett, F., Mendell, J.T., 2020. A ubiquitin ligase mediates target-directed microRNA decay independently of tailing and trimming. Science eabc9546. 10.1126/science.abc9546

Hatfield, S.D., Shcherbata, H.R., Fischer, K.A., Nakahara, K., Carthew, R.W., Ruohola-Baker, H., 2005. Stem cell division is regulated by the microRNA pathway. Nature 435, 974–978. 10.1038/nature03816

Hua, B.L., Orr-Weaver, T.L., 2017. DNA Replication Control During Drosophila Development: Insights into the Onset of S Phase, Replication Initiation, and Fork Progression. Genetics 207, 29–47. 10.1534/genetics.115.186627

Huang, Y.-C., Smith, L., Deng, W.-M., 2013. The microRNA miR-7 regulates Tramtrack69 in a developmental switch in Drosophila follicle cells. Development 140, 897–905. 10.1242/dev.080192

Iovino, N., Pane, A., Gaul, U., 2009. miR-184 has multiple roles in Drosophila female germline development. Developmental cell 17, 123–33. 10.1016/j.devcel.2009.06.008

Jackson, S.M., Blochlinger, K., 1997. cut interacts with Notch and protein kinase A to regulate egg chamber formation and to maintain germline cyst integrity during Drosophila oogenesis. Development 124, 3663–3672. 10.1242/dev.124.18.3663

Jiang, F., Ye, X., Liu, X., Fincher, L., McKearin, D., Liu, Q., 2005. Dicer-1 and R3D1-L catalyze microRNA maturation in Drosophila. Genes Dev. 19, 1674–1679. 10.1101/gad.1334005

Jin, Z., Xie, T., 2007. Dcr-1 Maintains Drosophila Ovarian Stem Cells. Current Biology 17, 539–544. 10.1016/j.cub.2007.01.050

Jones, B.T., Han, J., Zhang, H., Hammer, R.E., Evers, B.M., Rakheja, D., Acharya, A., Mendell, J.T., 2023. Target-directed microRNA degradation regulates developmental microRNA expression and embryonic growth in mammals. Genes Dev. 10.1101/gad.350906.123

Jordan, K.C., Schaeffer, V., Fischer, K.A., Gray, E.E., Ruohola-Baker, H., 2006. Notch signaling through Tramtrack bypasses the mitosis promoting activity of the JNK pathway in the mitotic-to-endocycle transition of Drosophila follicle cells. BMC Developmental Biology 6, 16. 10.1186/1471-213X-6-16

Khalisova, K.Y., Osadchiy, I.S., Georgiev, P.G., Maksimenko, O.G., 2021. TTK Isoforms Interact with Two Regions of the Mep-1 Protein of Drosophila melanogaster. Dokl Biochem Biophys 498, 177–179. 10.1134/S1607672921030042

Kingston, E.R., Bartel, D.P., 2021. Ago2 protects Drosophila siRNAs and microRNAs from target-directed degradation, even in the absence of 2′-O-methylation. RNA rna.078746.121. 10.1261/rna.078746.121

Kingston, E.R., Blodgett, L.W., Bartel, D.P., 2022. Endogenous transcripts direct microRNA degradation in Drosophila, and this targeted degradation is required for proper embryonic development. Molecular Cell S1097276522008498. 10.1016/j.molcel.2022.08.029

Kleaveland, B., Shi, C.Y., Stefano, J., Bartel, D.P., 2018. A Network of Noncoding Regulatory RNAs Acts in the Mammalian Brain. Cell 174, 350–362.e17. 10.1016/j.cell.2018.05.022

Knapp, E.M., Li, W., Sun, J., 2019. Downregulation of homeodomain protein Cut is essential for Drosophila follicle maturation and ovulation. Development 146, dev179002. 10.1242/dev.179002

König, A., Shcherbata, H.R., 2015. Soma influences GSC progeny differentiation via the cell adhesion-mediated steroid-let-7-Wingless signaling cascade that regulates chromatin dynamics. Biology Open 4, 285–300. 10.1242/bio.201410553

Kugler, J.-M., Chen, Y.-W., Weng, R., Cohen, S.M., 2013. miR-989 Is Required for Border Cell Migration in the Drosophila Ovary. PLOS ONE 8, e67075. 10.1371/journal.pone.0067075

Lee, S., Song, J., Kim, S., Kim, J., Hong, Y., Kim, Y., Kim, D., Baek, D., Ahn, K., 2013. Selective Degradation of Host MicroRNAs by an Intergenic HCMV Noncoding RNA Accelerates Virus Production. Cell Host & Microbe 13, 678– 690. 10.1016/j.chom.2013.05.007

Levine, B., Hackney, J.F., Bergen, A., Dobens, L., Truesdale, A., Dobens, L., 2010. Opposing interactions between Drosophila Cut and the C/EBP encoded by Slow Border Cells direct apical constriction and epithelial invagination. Developmental Biology 344, 196–209. 10.1016/j.ydbio.2010.04.030

Li, L., Sheng, P., Li, T., Fields, C.J., Hiers, N.M., Wang, Y., Li, J., Guardia, C.M., Licht, J.D., Xie, M., 2021. Widespread microRNA degradation elements in target mRNAs can assist the encoded proteins. Genes Dev. 35, 1595–1609. 10.1101/gad.348874.121

Libri, V., Helwak, A., Miesen, P., Santhakumar, D., Borger, J.G., Kudla, G., Grey, F., Tollervey, D., Buck, A.H., 2012. Murine cytomegalovirus encodes a miR-27 inhibitor disguised as a target. PNAS 109, 279–284. 10.1073/pnas.1114204109

López-Schier, H., Johnston, D.S., 2001. Delta signaling from the germ line controls the proliferation and differentiation of the somatic follicle cells during Drosophila oogenesis. Genes & development 15, 1393–1405. 10.1101/gad.200901

Marcinowski, L., Tanguy, M., Krmpotic, A., Rädle, B., Lisnić, V.J., Tuddenham, L., Chane-Woon-Ming, B., Ruzsics, Z., Erhard, F., Benkartek, C., Babic, M., Zimmer, R., Trgovcich, J., Koszinowski, U.H., Jonjic, S., Pfeffer, S., Dölken, L., 2012. Degradation of Cellular miR-27 by a Novel, Highly Abundant Viral Transcript Is Important for Efficient Virus Replication In Vivo. PLOS Pathogens 8, e1002510. 10.1371/journal.ppat.1002510 Marco, A., 2015. Selection Against Maternal microRNA Target Sites in Maternal

Transcripts. G3: Genes, Genomes, Genetics 5, 2199–2207. 10.1534/g3.115.019497

Marco, A., 2014. Sex-biased expression of microRNAs in Drosophila melanogaster. Open biology 4, 140024. 10.1098/rsob.140024

Montell, D.J., 2003. Border-cell migration: the race is on. Nat Rev Mol Cell Biol 4, 13–24. 10.1038/nrm1006

Montell, D.J., Rorth, P., Spradling, A.C., 1992. slow border cells, a locus required for a developmentally regulated cell migration during oogenesis, encodes Drosophila CEBP. Cell 71, 51–62. 10.1016/0092-8674(92)90265-E

Nahar, S., Morales Moya, L.J., Brunner, J., Hendriks, G.-J., Towbin, B., Hauser, Y.P., Brancati, G., Gaidatzis, D., Großhans, H., 2024. Dynamics of miRNA accumulation during C. elegans larval development. Nucleic Acids Research 52, 5336–5355. 10.1093/nar/gkae115

Nakahara, K., Kim, K., Sciulli, C., Dowd, S.R., Minden, J.S., Carthew, R.W., 2005. Targets of microRNA regulation in the *Drosophila* oocyte proteome. Proc. Natl. Acad. Sci. U.S.A. 102, 12023–12028. 10.1073/pnas.0500053102

Ni, J.-Q., Zhou, R., Czech, B., Liu, L.-P., Holderbaum, L., Yang-Zhou, D., Shim, H.- S., Tao, R., Handler, D., Karpowicz, P., Binari, R., Booker, M., Brennecke, J., Perkins, L.A., Hannon, G.J., Perrimon, N., 2011. A genome-scale shRNA resource for transgenic RNAi in Drosophila. Nat Methods 8, 405–407. 10.1038/nmeth.1592

Ogienko, A.A., Yarinich, L.A., Fedorova, E.V., Lebedev, M.O., Andreyeva, E.N., Pindyurin, A.V., Baricheva, E.M., 2018. New slbo-Gal4 driver lines for the analysis of border cell migration during Drosophila oogenesis. Chromosoma 127, 475–487. 10.1007/s00412-018-0676-7

Papantonis, A., Swevers, L., Iatrou, K., 2015. Chorion Genes: A Landscape of Their Evolution, Structure, and Regulation. Annu. Rev. Entomol. 60, 177–194. 10.1146/annurev-ento-010814-020810

Park, J.K., Liu, X., Strauss, T.J., McKearin, D.M., Liu, Q., 2007. The miRNA Pathway Intrinsically Controls Self-Renewal of Drosophila Germline Stem Cells. Current Biology 17, 533–538. 10.1016/j.cub.2007.01.060

Poulton, J.S., Huang, Y.-C., Smith, L., Sun, J., Leake, N., Schleede, J., Stevens, L.M., Deng, W.-M., 2011. The microRNA pathway regulates the temporal pattern of Notch signaling in *Drosophila* follicle cells. Development 138, 1737–1745. 10.1242/dev.059352

Rauluseviciute, I., Riudavets-Puig, R., Blanc-Mathieu, R., Castro-Mondragon, J.A., Ferenc, K., Kumar, V., Lemma, R.B., Lucas, J., Chèneby, J., Baranasic, D., Khan, A., Fornes, O., Gundersen, S., Johansen, M., Hovig, E., Lenhard, B., Sandelin, A., Wasserman, W.W., Parcy, F., Mathelier, A., 2024. JASPAR 2024: 20th anniversary of the open-access database of transcription factor binding profiles. Nucleic Acids Research 52, D174–D182. 10.1093/nar/gkad1059

Rørth, P., Szabo, K., Texido, G., 2000. The Level of C/EBP Protein Is Critical for Cell Migration during Drosophila Oogenesis and Is Tightly Controlled by Regulated Degradation. Molecular Cell 6, 23–30. 10.1016/S1097-2765(05)00008-0

Saadin, A., Starz-Gaiano, M., 2016. Circuitous Genetic Regulation Governs a Straightforward Cell Migration. Trends in Genetics 32, 660–673. 10.1016/j.tig.2016.08.001

Shcherbata, H.R., Ward, E.J., Fischer, K.A., Yu, J.-Y., Reynolds, S.H., Chen, C.-H., Xu, P., Hay, B.A., Ruohola-Baker, H., 2007. Stage-Specific Differences in the Requirements for Germline Stem Cell Maintenance in the Drosophila Ovary. Cell Stem Cell 1, 698–709. 10.1016/j.stem.2007.11.007

Sheng, P., Li, L., Li, T., Wang, Y., Hiers, N.M., Mejia, J.S., Sanchez, J.S., Zhou, L., Xie, M., 2023. Screening of Drosophila microRNA-degradation sequences reveals Argonaute1 mRNA’s role in regulating miR-999. Nat Commun 14, 2108. 10.1038/s41467-023-37819-9

Sheu-Gruttadauria, J., Pawlica, P., Klum, S.M., Wang, S., Yario, T.A., Schirle Oakdale, N.T., Steitz, J.A., MacRae, I.J., 2019. Structural Basis for Target-Directed MicroRNA Degradation. Molecular Cell 75, 1243–1255.e7. 10.1016/j.molcel.2019.06.019

Shi, C.Y., Elcavage, L.E., Chivukula, R.R., Stefano, J., Kleaveland, B., Bartel, D.P., 2023. ZSWIM8 destabilizes many murine microRNAs and is required for proper embryonic growth and development. Genome Res. gr.278073.123. 10.1101/gr.278073.123

Shi, C.Y., Kingston, E.R., Kleaveland, B., Lin, D.H., Stubna, M.W., Bartel, D.P., 2020. The ZSWIM8 ubiquitin ligase mediates target-directed microRNA degradation. Science 370. 10.1126/science.abc9359

Simeone, I., Rubolino, C., Noviello, T.M.R., Farinello, D., Cerulo, L., Marzi, M.J., Nicassio, F., 2022. Prediction and pan-cancer analysis of mammalian transcripts involved in target directed miRNA degradation. Nucleic Acids Research 50, 2019–2035. 10.1093/nar/gkac057

Sokol, N.S., Xu, P., Jan, Y.-N., Ambros, V., 2008. *Drosophila let-7* microRNA is required for remodeling of the neuromusculature during metamorphosis. Genes Dev. 22, 1591–1596. 10.1101/gad.1671708

Stubna, M.W., Shukla, A., Bartel, D.P., 2024. Widespread destabilization of C. elegans microRNAs by the E3 ubiquitin ligase EBAX-1. RNA rna.080276.124. 10.1261/rna.080276.124

Sun, J., Deng, W.-M., 2007. Hindsight Mediates the Role of Notch in Suppressing Hedgehog Signaling and Cell Proliferation. Developmental Cell 12, 431–442. 10.1016/j.devcel.2007.02.003

Sun, J., Deng, W.-M., 2005. Notch-dependent downregulation of the homeodomain gene cut is required for the mitotic cycle/endocycle switch and cell differentiation in *Drosophila* follicle cells. Development 132, 4299–4308. 10.1242/dev.02015

Sun, J., Smith, L., Armento, A., Deng, W.-M., 2008. Regulation of the endocycle/gene amplification switch by Notch and ecdysone signaling. Journal of Cell Biology 182, 885–896. 10.1083/jcb.200802084

Vilmos, P., Bujna, Á., Szuperák, M., Havelda, Z., Várallyay, É., Szabad, J., Kucerova, L., Somogyi, K., Kristó, I., Lukácsovich, T., Jankovics, F., Henn, L., Erdélyi, M., 2013. Viability, Longevity, and Egg Production of Drosophila melanogaster Are Regulated by the miR-282 microRNA. Genetics 195, 469–480. 10.1534/genetics.113.153585

Wang, C., Jia, Q., Guo, X., Li, K., Chen, W., Shen, Q., Xu, C., Fu, Y., 2022. microRNA-34 family: From mechanism to potential applications. The International Journal of Biochemistry & Cell Biology 144, 106168. 10.1016/j.biocel.2022.106168

Wang, G., Lei, J., Wang, Y., Yu, J., He, Y., Zhao, W., Hu, Z., Xu, Z., Jin, Y., Gu, Y., Guo, X., Yang, B., Gao, Z., Wang, Z., 2023. The ZSWIM8 ubiquitin ligase regulates neurodevelopment by guarding the protein quality of intrinsically disordered Dab1. Cerebral Cortex 33, 3866–3881. 10.1093/cercor/bhac313

Wu, X., Tanwar, P.S., Raftery, L.A., 2008. Drosophila follicle cells: Morphogenesis in an eggshell. Seminars in Cell & Developmental Biology, Cell Shape and Tissue Morphogenesis 19, 271–282. 10.1016/j.semcdb.2008.01.004

Xia, J., Wang, L., Lei, F., Pan, L., Liu, L., Wan, P., 2024. MicroRNA-34 disrupts border cell migration by targeting *Eip74EF* in *Drosophila melanogaster*. Journal of Insect Physiology 104724. 10.1016/j.jinsphys.2024.104724

Yang, L., Chen, Dongsheng, Duan, R., Xia, L., Wang, J., Qurashi, A., Jin, P., Chen, Dahua, 2007. Argonaute 1 regulates the fate of germline stem cells in Drosophila. Development (Cambridge, England) 134, 4265–72. 10.1242/dev.009159

Yatsenko, A.S., Shcherbata, H.R., 2018. Stereotypical architecture of the stem cell niche is spatiotemporally established by miR-125-dependent coordination of Notch and steroid signaling. Development dev.159178. 10.1242/dev.159178

Yoon, W.H., Meinhardt, H., Montell, D.J., 2011. miRNA-mediated feedback inhibition of JAK/STAT morphogen signalling establishes a cell fate threshold. Nat Cell Biol 13, 1062–1069. 10.1038/ncb2316

Yu, J.-Y., Reynolds, S.H., Hatfield, S.D., Shcherbata, H.R., Fischer, K.A., Ward, E.J., Long, D., Ding, Y., Ruohola-Baker, H., 2009. Dicer-1-dependent Dacapo suppression acts downstream of Insulin receptor in regulating cell division of Drosophila germline stem cells. Development 136, 1497–1507. 10.1242/dev.025999

